# Genomic adaptation in the CAZyome and specialised metabolism of the plant-associated *Streptomyces violaceusniger* clade

**DOI:** 10.1101/2021.10.25.465742

**Authors:** Damien Gayrard, Marine Veyssière, Clément Nicolle, Kévin Adam, Yves Martinez, Céline Vandecasteele, Marie Vidal, Bernard Dumas, Thomas Rey

## Abstract

Streptomycetes are Gram-positive actinobacteria largely represented in the plant root microbiota. The genetic determinants involved in the presence of *Streptomyces* in the rhizosphere are mostly unknown but can rely on the ability to release phytohormones, degrade plant cell-wall polysaccharides and produce specialised metabolites. Here we sequenced the genome of the rhizospheric and plant defence-stimulating strain *Streptomyces* sp. AgN23. We found out that it belongs to the soil and plant root dwelling *S. violaceusniger* clade. The genome annotation of AgN23 revealed the ability of the bacterium to synthesise auxin, a major regulator of plant development, to degrade plant cell wall with a large repertoire of carbohydrate degrading enzymes and to produce antimicrobials (rustmicin, mediomycin, niphimycin, nigericin) and plant bioactive compounds (nigericin, echosides, elaiophylin) through a set of biosynthetic gene clusters. We also found that these genomic features are well-conserved among members of the *S. violaceusniger* clade. In addition, AgN23 display original events of biosynthetic gene clusters acquisitions and losses which may account for its beneficial effect on plants. Taken together, our work supports the hypothesis that hydrolytic enzymes and specialised metabolites repertoires underpin the interaction of bacteria belonging to the *S. violaceusniger* clade with plant roots within the rhizosphere.

**Impact statement:** Streptomycetes are filamentous Gram-positive bacteria universally found around and within host plant tissues. These actinobacteria have been extensively investigated for their tremendous ability to produce diverse specialised metabolites (e.g., antibiotics). By contrast their impact on host plant physiology is widely neglected. Whether specific lineage of *Streptomyces* colonise host plant and what are the underlying molecular mechanisms is poorly documented. Here we report a chromosome-scale assembly of AgN23 genome, a *Streptomyces* sp. strain previously characterised for its ability to activate the plant immune system. This reference sequence enabled us to position AgN23 in the *S. violaceusniger* clade from which several representatives have been isolated worldwide from the rhizosphere of unrelated plants. Comparative genomic studies suggest that *S. violaceusniger* spp. produce a prominent CAZyome with expansion of plant cell wall degrading enzymes families and a conserved specialised metabolism acting on host plant physiology and its rhizospheric microbiota. These genomic features may underly *S. violaceusniger* spp. adaptation to the rhizopsheric niche.

**Data summary:** The raw reads sequences of AgN23 genome are available at NCBI on the Sequence Read Archive portal for PacBio and MiSeq data (SRR13990229 and SRR14028548 respectively). The Genome assembly is available on the NCBI nucleotide portal under the accession NZ_CP007153.1. This genome sequence was uploaded on the MicroScope platform for genome annotation and analysis (https://mage.genoscope.cns.fr/microscope/home/index.php) [1]. The RNA-seq raw reads are archived in the NCBI Bioproject PRJNA745930. The following eight supplementary tables are included in the online version of this article.

Supplementary Information 1: Genomes used in this study. The accession number used from the NCBI portal, name, size, number of contigs as well as the level of completeness of the assembly are indicated.

Supplementary Information 2: List of the single copy core genes used by autoMLST to build the phylogenetic tree in Figure 1.

Supplementary information 3: Annotation of AgN23 full chromosome. For each gene the frame of translation, sequence length and position on the chromosome are indicated. All genes were annotated according to the Microscope platform, see materials and methods. In addition, the expression for each gene is reported in transcripts per million (TPM) based on the the RNA-seq data from three biological replicates.

Supplementary Information 4: Genomes having a Mash-based estimated ANI (Average Nucleotide Identity) superior or egal to 80% according to autoMLST.

Supplementary Information 5: Prediction of the CAZyme encoding genes using HMMER dbCAN2. The genes are sorted according their CAZy families. For each gene, the begin position on the chromosome, the CAZy category, the annotation, the expression level in transcripts per million (TPM) and the predicted targets of the putative enzymes are described.

Supplementary Information 6: Gene identified by antiSMASH in the region containing a biosynthetic gene cluster. Expression levels in transcripts per million (TPM) are indicated for each gene. Annotated central bioynthetic genes are indicated as Y. Those are the ones used for the calculation of mean BGC expression in Table 2.

Supplementary Information 7: Annotation of AgN23 genes putatively involved in biosynthetic pathways for Auxins related phytohomones. Expression levels in transcripts per million (TPM) are indicated for each gene. The genes were detected by blasting reference KEGG sequences for each KEGG ONTOLOGY against AgN23 genes. A cut off of 70% identity and 40% coverage was applied to detect positive hits. These biosynthetic pathways and the KEGG ONTOLOGY are indicated in column F and G.

Supplementary Information 8: Inspection of BiG-FAM hits with AgN23 BGCs to identify homologous BGCs found outside the S. violaceusniger clade. BiG-FAM distance higher than 900 were excluded from the analysis.

**The authors confirm all supporting data, code and protocols have been provided within the article or through supplementary data files**.

## Introduction

Streptomycetes are aerobic and Gram-positive actinobacteria forming branched vegetative mycelium before developing aerial hyphae bearing spores [2]. These bacteria received considerable attention from a biotechnological point of view, notably regarding their enzymatic repertoire [3] and in the drug discovery field, leading to the structure elucidation of more than 6000 specialised metabolites [4, 5]. Their tremendous ability to produce antimicrobial compounds relies on the wealth and diversity of Biosynthetic Gene Clusters (BGCs) encoded in their genomes [6]. Since several of the metabolites they produce display strong antimicrobial activities, *Streptomyces* recruitment in the rhizosphere and inside root tissues presumably protect the host from pathogens. The recent soaring of microbial metabarcoding approaches have highlighted *Streptomyces* spp. prominent abundance in plant roots microbiota [7, 8]. *Streptomyces* based products have been developed for agriculture, but these strains only cover a subset of plant colonising *Streptomyces* species [9, 10]. Notwithstanding, little is known regarding the biological function of these bacteria in the plant environment and the gene families involved in their adaptation to this ecological niche [11, 12]. More than 100 hundred fully assembled genome sequences of *Streptomyces* strains are currently available, paving the way to genome-based phylogeny and comparisons of BGCs content across *Streptomyces* clades [13-15]. This knowledge open inroads to rationalise and investigate the potential use of *Streptomyces* strains in agriculture.

Here we report a gapless assembly of *Streptomyces* sp. AgN23 (AgN23), previously isolated from grapevine rhizosphere and identified as a strong inducer of plant defences [16]. We used this high-quality assembly to position AgN23 in the *Streptomyces violaceusniger* genomospecies. We then dissected the original features in the CAZyome and BGC content of AgN23 and other *Streptomyces violaceusniger* strains and shed light on specificities in the specialised metabolism and carbohydrate degradation capabilities of this lineage. This comparative genomic approach also lead us to identify original feature in the genome of AgN23 such as putative BGCs losses and acquisition by horizontal gene transfer which may account for the bacteria interaction with plants.

## Methods

### AgN23 cultivation and HMW DNA extraction

AgN23 strain was cultivated as described previously [16, 17]. In brief the strain was grown on solid modified Bennet medium (D-Glucose 10 g/l; Soybean peptones 2.5 g/l; Yeast Extract 1.5 g/l; Agar 16 g/l) or International Streptomyces Project media ISP2, ISP3, ISP4 and ISP5 [18, 19]. To produce the spores inoculum, we incubated Bennet plates for two weeks at 22°C in the darkness before filling them with 10 ml of sterile water. The mycelium was scraped with a spreader and the resulting solution was filtered in 50 ml syringe filled with non-absorbent cotton wool. For DNA extraction the AgN23 mycelium was grown at 28°C and under 250 rpm in 250 ml Erlenmeyer flasks containing 50 ml of liquid Bennet. Approximately 100 mg of AgN23 pellets were collected by centrifugation at 11 000 g and flash frozen in liquid nitrogen. Genomic DNA was isolated using the Nucleobond RNA/DNA kit (Macherey-Nagel) according to the manufacturer’s instructions.

### Library preparation for genome sequencing

Library preparation and sequencing were performed at the GeT-PlaGe core facility (Castanet-Tolosan), according to the PacBio’s instructions with 15 kb size cut-off (“20 kb Template Preparation Using BluePippin™ Size Selection system”). At each step, DNA was quantified using the Qubit dsDNA HS Assay Kit (Life Technologies). DNA purity was tested using the NanoDrop (Thermo Fisher) and size distribution and degradation assessed using High Sensitivity Large Fragment 50kb Analysis Kit used with a Fragment analyser (AATI). Purification steps were performed using 0.45X AMPure PB beads (PacBio). A total of 10 µg of DNA was purified then sheared at 40kb using the Megaruptor system (Diagenode). The Single Molecule Real-Time (SMRT) sequencing was performed using SMRTBell template Prep Kit 1.0 (PacBio), a DNA and END damage repair step was performed on 5 µg of sample. Then, blunt hairpin adapters were ligated to the library. The library was treated with an exonuclease cocktail to digest unligated DNA fragments. A size selection step using a 10kb cut-off was performed on the BluePippin Size Selection system (Sage Science) with 0.75% agarose cassettes, Marker S1 high Pass 15-20 kb. Conditioned Sequencing Primer V2 was annealed to the size selected SMRTbell. The annealed library was then bound to the P6-C4 polymerase using a ratio of polymerase to SMRTbell at 10:1. Then after a magnetic bead-loading step (OCPW), SMRTcell libraries were sequenced on 2 SMRTcells on RSII instrument at 0.18 to 0.23 nM with a 360 min movie. The initially generated raw sequencing reads were evaluated in terms of the average quality score at each position, GC content distribution, quality distribution, base composition, and other metrics. Sequencing reads with low quality were also filtered out before the genome assembly and annotation of gene structure. Finally, microbial DNA potential contamination was excluded after comparison by BLAST of the draft assembly of the first SMRT cell against a 16S ribosomal RNA sequences data bank (Bacteria and Archaea).

### Genome assembly, annotation, and comparative genomics

The subreads were assembled with PacBio’s SMRT analysis software version 2.3.0 using default settings with a minimum subreads length of 3 kb to exclude smaller sequenced reads and a read score of better than 0.8 to enrich in reads with a low error rate. The single unitig obtained by long-read sequencing was corrected with Mi-Seq (Illumina®) data using Pilon (version 1.21), resulting in 165 substitutions and two deletions of 44 and 5 bases. This final genome assembly was retained for subsequent analysis. The gene annotation was performed on the MicroScope Microbial Genome Annotation & Analysis Platform [1]. Genome completion was checked with CheckM and BUSCO [20, 21]. CAZy genes annotation was obtained running the HMMER tool (e-value < 1e-15, coverage > 0.35) on the dbCAN2 meta server [22]. For the annotation of auxin biosynthesis genes, we retrieved the protein sequences from UniProt corresponding to the KEGG ontologies associated to the different biosynthetic steps described in Zhang *et al*. [23]. These sequences were systematically used as query for BLASTx analysis against AgN23 chromosome sequences. The position on AgN23 chromosome of any hits showing more than 70% identity and 40% coverage with one query was used to determine the corresponsing AgN23 gene model [23]. antiSMASH 6.0 was used to detect BGC-containing regions in the AgN23 chromosome and annotate detected sequences based on the MIBiG 2.0 repository [24, 25]. The comparative genomics was performed with BiG-SCAPE using default parameters with antiSMASH 6.0 predicted region-containing BGCs as input data to produce similarity networks [26]. Complementarily, the BiG-SLiCE software (1.0.0) was used to identify BGCs showing similarity to AgN23 regions annotated by antiSMASH among the 1,225,071 BGCs stored in the BiG-FAM database [15, 27]. BiG-SLICE was used with standard parameters, including the arbitrary clustering threshold (T=900.0).

### Phylogenetic and phylogenomic analysis

The phylogenetic analysis of *Streptomyces* sp. AgN23 was performed using genome sequences previously assignated to the *S. violaceusniger* clade (*Streptomyces* sp. M56, *Streptomyces* sp. 11-1-2, *S. rapamycinicus* NRRL 5491, S. malaysiensis DSM4137, S. *antimycoticus* NBRC 100767, S. *sabulosicollis* PRKS01-29, S. *albiflaviniger* DSM 41598, S. *rhizosphaericus* NRRL B-24304, S. *javensis* DSM 41764 and S. *milbemycinicus* NRRL) [28-32]. In addition, we introduced as input the genomes of the model strain S. *coelicolor* A(3)2 and of two biocontrol strains, S. *lydicus* WYEC108 and S. *griseoveridis* K61 as well as *Frankia alni* ACN14a as outgroup (Supplementary Information 1). The *Streptomyces* phylogeny was built using a Multi-Locus Sequence Typing (MLST) strategy with the *de* novo workflow of autoMLST [33]. The concatenated alignment of 85 single copy conserved genes was built using the Fast alignment mode (MAFFT FFT-NS-2) and the IQ-TREE Ultrafast Bootstrap analysis was performed with 1000 replicates (Supplementary Information 2)[33].

Finally, a whole-genome based phylogenetic tree based on fully-assembled chromosome of isolates from the *S. violaceusniger* clade has been inferred with FastME 2.1.6.1 using distance calculated by GBDP (Genome Blast Distance Phylogeny) available on the TYGS platform [34, 35]. The branch lengths were scaled in terms of GBDP distance formula d5. The tree had an average branch support of 96.9% after 100 bootstrap replicates and was rooted at midpoint.

### Library preparation for transcriptome sequencing and expression analysis

AgN23 was cultivated in liquid Bennett medium at 28°C for 48h. Total RNA was isolated using the RNeasy Plant Mini Kit (Qiagen) according to manufacturer’s instructions. Three replicates were prepared for the construction of the libraries. rRNA depletion was performed using Ribozero rRNA Removal Kit (Illumina). RNA sequencing library preparation used NEBNext Ultra RNA Library Prep Kit for Illumina following the manufacturer’s recommendations (NEB). RNAs were fragmented to generate double stranded cDNA, subsequently ligated to universal adapter, followed by index addition and library enrichment with limited cycle PCR. Sequencing libraries were validated using the Agilent Tapestation 4200 (Agilent Technologies) and quantified by Qubit 2.0 Fluorometer (Invitrogen) as well as quantitative PCR (Applied Biosystems). RNA-Seq experiments have been performed on an Illumina HiSeq4000 using a paired-end read length of 2×150 bp with the Illumina HiSeq4000 sequencing kits. The mapping and statistical analysis was performed using the bioinformatics pipeline implemented in the MicroScope Platform [1]. Raw reads of each sample (R1 fastq files from the paired-end run) were mapped onto *Streptomyces* sp. AgN23 reference genome with BWA-MEM (v.0.7.4) [36]. An alignment score equal to at least half of the read was required for a hit to be retained. SAMtools (v.0.1.8) was then used to extract reliable alignments with a Mapping Quality (MAPQ)>=1 from SAM formatted files [37]. The number of reads matching each genomic object of the reference sequence was subsequently counted with the toolset BEDTools (v.2.10.1) [38]. The mean transcripts per million (TPM) for each gene was then calculated from the three independent samples. The BGCs expression levels have been determined by calculating the mean expression of central biosynthetic genes for each BGC as defined by antiSMASH. A mean expression for each BGC across the three biological repetitions was then determined.

### Scanning electronic microscopy

The observation of AgN23 mycelium and spore development by Scanning Electronic microscopy was performed on a Quanta 250 FEG (FEI). Agar plugs of two weeks old AgN23 cultures were placed on micrometric platen, frozen in liquid nitrogen and finally metallized with platinum. The samples were observed microscopically at an accelerating voltage of 5.00 kV.

## Results and discussion

### Chromosome-scale assembly of AgN23 and genome-based taxonomic assignation to the S. *violaceusniger* clade

We performed a PacBio® RSII long-read sequencing and obtained a linear chromosome of 10,86 Mb for AgN23. This genome sequence was polished using Illumina® MiSeq sequences to produce the final assembly. The final assembly was uploaded on the MicroScope Platform [1]. The genome annotation retrieved 10,514 protein coding sequences, 10,458 of them being supported by RNA-seq with expression levels comprised from 0.09 to 20661 transcripts per million (TPM) and mean and median values of 94 and 21 TPM respectively (Supplementary Information 3)(Table1). A checkM analysis was performed using 455 genomes and 315 lineage-specific markers and has validated the completeness of the assembly and the annotation (Table 1)[20]. Complementarly, the completeness was also supported by a BUSCO analysis that detected 99.7% of the genes that are expected in the genome of an actinobacterium (Table 1)[21].

**Table 1:** Summary of the assembly and the annotation of *Streptomyces* sp. AgN23 complete chromosome obtained by PacBio® and Illumina® sequencing.

| **Assembly Statistics** | ***Streptomyces* sp. AgN23** |
| --- | --- |
| **Genome size (bp)** | 10,858,739 |
| **Coverage (PacBio®)** | 77X |
| **G+C content (%)** | 70.9 |
| **Number of protein coding genes** | 10,514 |
| **CheckM Completeness** | 100% |
| **Contamination** | 1.84% |
| **BUSCO** | 451/452 |
| **CAZymes** | 278 |
| **Number BGCs (antiSMASH 6.0)** | 47 |

More than 500 *Streptomyces* species have been described based on their 16S rRNA sequence. Previous sequencing of AgN23 16S rRNA showed strongest conservation with representatives of the *S. violaceusniger* clade, most notably *Streptomyces castelarensis* [16]. However, lack of variation in 16S rRNA may confound strains belonging to different species [39]. Recent development of long-read technologies and massive sequencing of *Streptomyces* has leveraged genome based phylogenies [6]. Thus, we decided to consolidate the taxonomic affiliation of AgN23 with the gapless assembly of AgN23 chromosome. We performed an autoMLST approach based on publicly available *Streptomyces* sequences. In brief, 85 conserved housekeeping genes showing neutral dN/dS were concatenated and aligned as a basis for tree building (Figure 1, Supplementary Information 2) [33]. As a result, using AgN23 as query, we affiliated the strain to a clade containing six other strains showing an Average Nucleotide Identity (ANI) higher than 95% with AgN23 which is conventionally considered as a threshold for species delimitation [40]. It contains the closely related *S. melanosporofaciens* DSM 40318, *S. antimycoticus* NBRC 100767, *S. violaceusniger* NRRL F-8817, *S. violaceusniger* Tu 4113, *S. hygroscopicus* XM201 and *Streptomyces* sp. 11-1-2 [41, 42]. In total, 28 isolates harboured ANI>85% with *Streptomyces* sp. AgN23. Noteworthy, this group contains hallmark representative species of the *S. violaceusniger* clade, such as, *S. hygroscopicus, S. sparsogenes, S. malaysiensis* [43], *S. himastatinicus, S. rapamycinicus*, and other close species *S. autolyticus* [44], *S. antioxidans* [45] and *S. iranensis* [46] *S. albiflaviniger, S. rhizosphaericus, S*. sabulosicollis, *S. javensis* and *S. milbemycinicus* [30](Supplementary Information 4). Interestingly, the strains *Streptomyces* strains RT-d22 [47], *Streptomyces* sp. Strain PRh5 [48], *Streptomyces* sp. 11-1-2 [29, 49], *S. hygroscopicus* Osish-2 [50-54], *Streptomyces* sp. NBRC 109436 [55] and *S. rhizosphaericus* NRRL B-24304 [32] were isolated from the rhizosphere of diverse plants across the world. This suggests that interaction with plants is widespread among the strains belonging to the *S. violaceusniger* clade. Finally, cultivation of AgN23 on a range of ISP media confirmed that AgN23 displays typical phenotypes of S. *violaceusniger* clade with whitish colony then turning grey during the sporulation process and resulting in the formation of spiralled chains of rugose-ornamented spores (Figure 2)[31, 32, 56].

**Figure 1:**
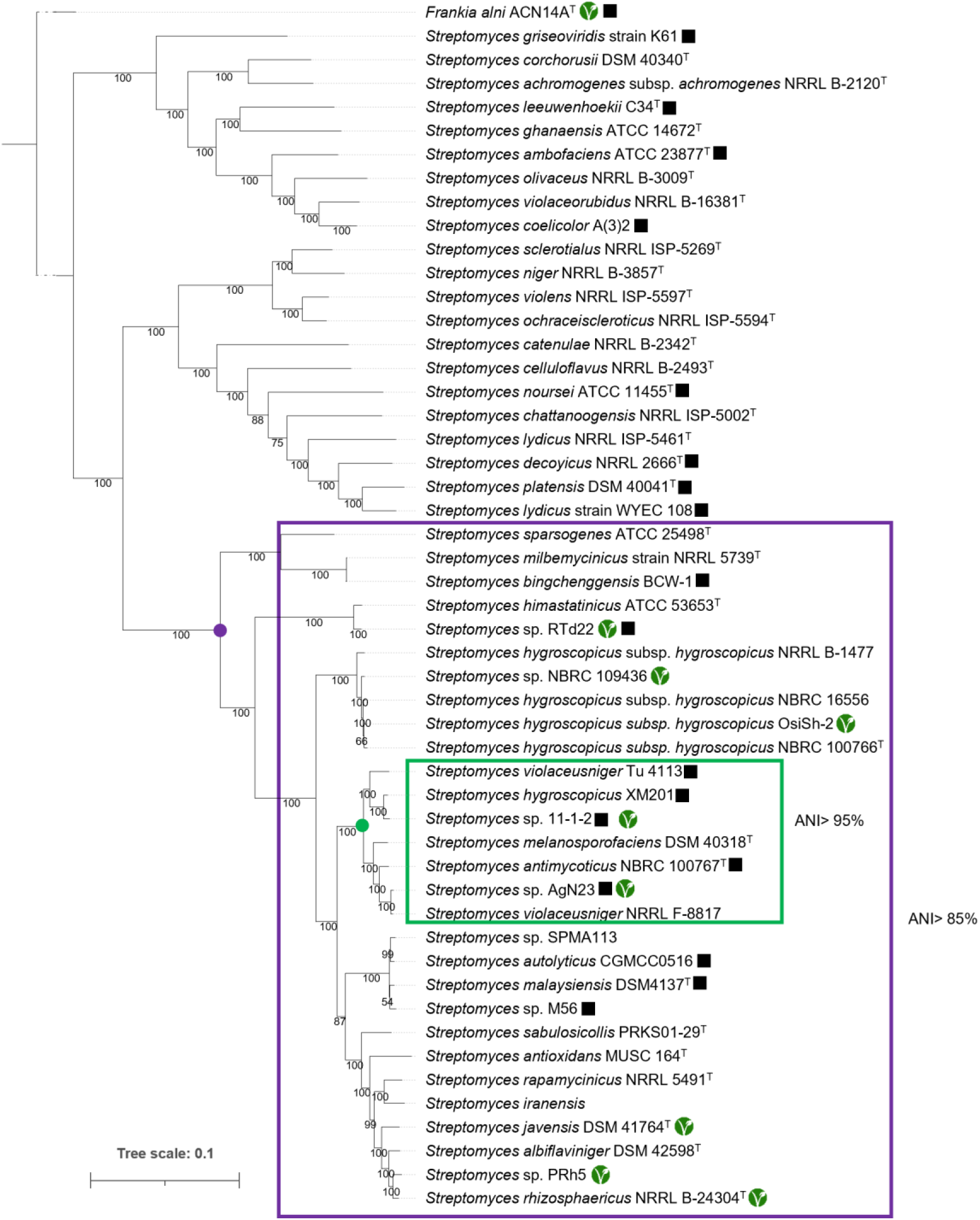
Multi-locus sequence typing assigned AgN23 to the *S. violaceusniger* clade. Phylogenetic tree based on the multiple alignment of 85 single-copy homologous genes selected from genomic sequences with AutoMLST. The green node highlights the isolates considered to be from the same species (ANI>95%). The black node highlights the clade formed by isolates with ANI>90% as compared to AgN23. The black squares highlights the eight strains that were used for the BGC conservation study. The green logo indicates plant-isolated strains. *Frankia alni* ACN14a was used as outgroup, bootstrap=100.

**Figure 2:**
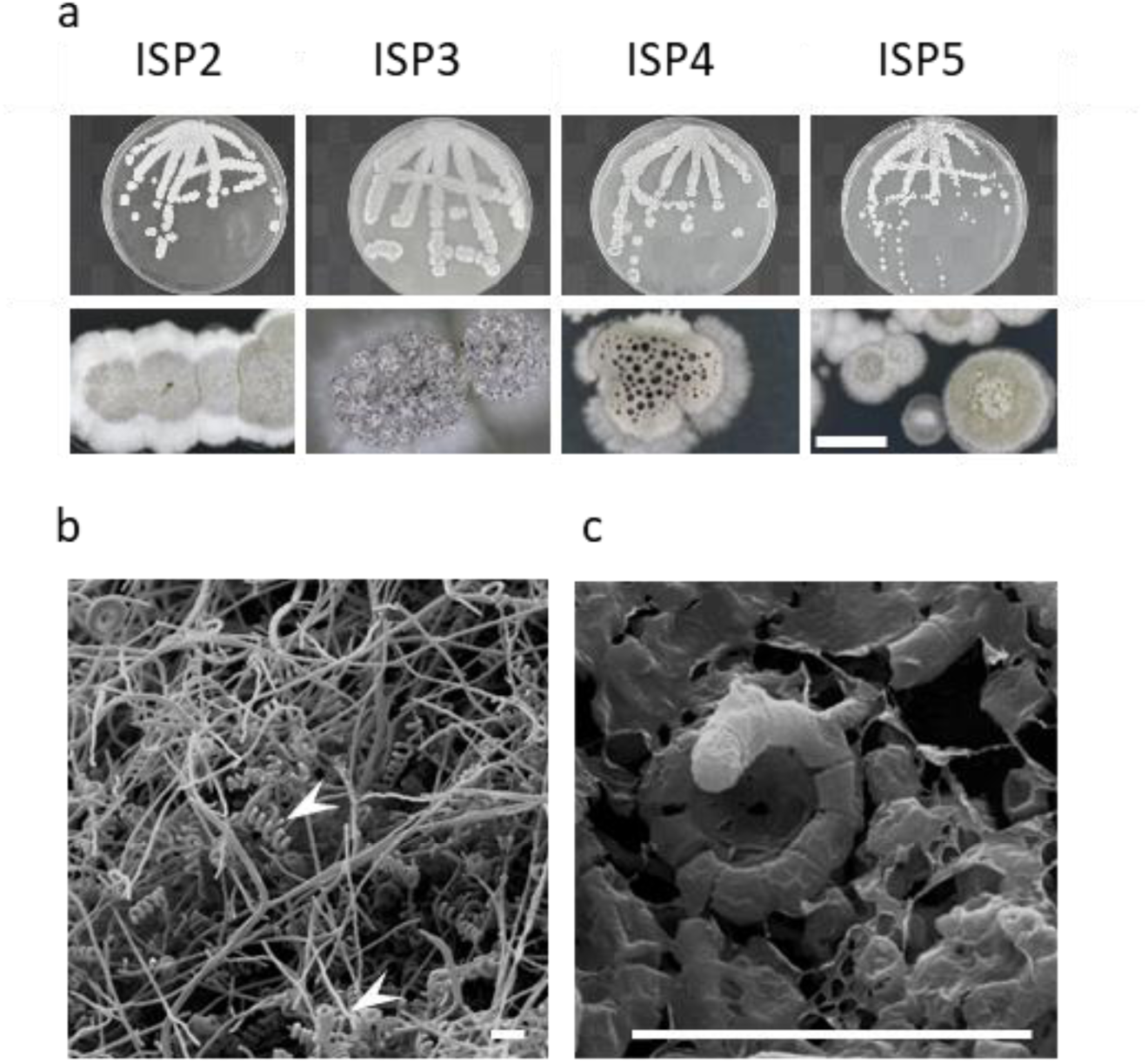
AgN23 harbours typical phenotypic features of the *S. violaceusniger* clade. a) AgN23 forms a white mycelium turning grey at the onset of sporulation as observed on a range of ISP media, the Petri plate are 9 cm in diameter and the scale bar is 1mm. b) Scanning Electron Microscopy observation of spiralled chains of spores (white arrow) are formed by AgN23 (scale bar = 6µm). c) Scanning Electron Microscopy observation of AgN23 spores showing a rugose-ornamented surface (scale bar = 6µm).

### AgN23 exhibits a wide repertoire of CAZymes related to plant cell wall degradation

Streptomycetes display extensive abilities to degrade polysaccharides based on their rich repertoires of Carbohydrate-Active Enzymes (CAZymes) [3, 22]. We annotated the AgN23 CAZyome with dbCAN2 and found 278 CAZymes representing 2.6% of AgN23 coding sequences (Supplementary Information 5). The *Streptomyces* CAZyome typically ranges from 1.5% to 3% of the proteome [3]. For example, CAZymes represents 2.9% of coding sequences of the hallmark cellulolytic *Streptomyces* sp. SirexAA-E, thus AgN23 genome harbours a rather large CAZyome [3]. To investigate whether AgN23 CAZyome displays expanded repertoire in particular CAZy families, we compared it with the CAZyome of *Streptomyces* sp. SirexAA-E and the 50 genomes used in the phylogenomic analysis (Figure 3).

**Figure 3:**
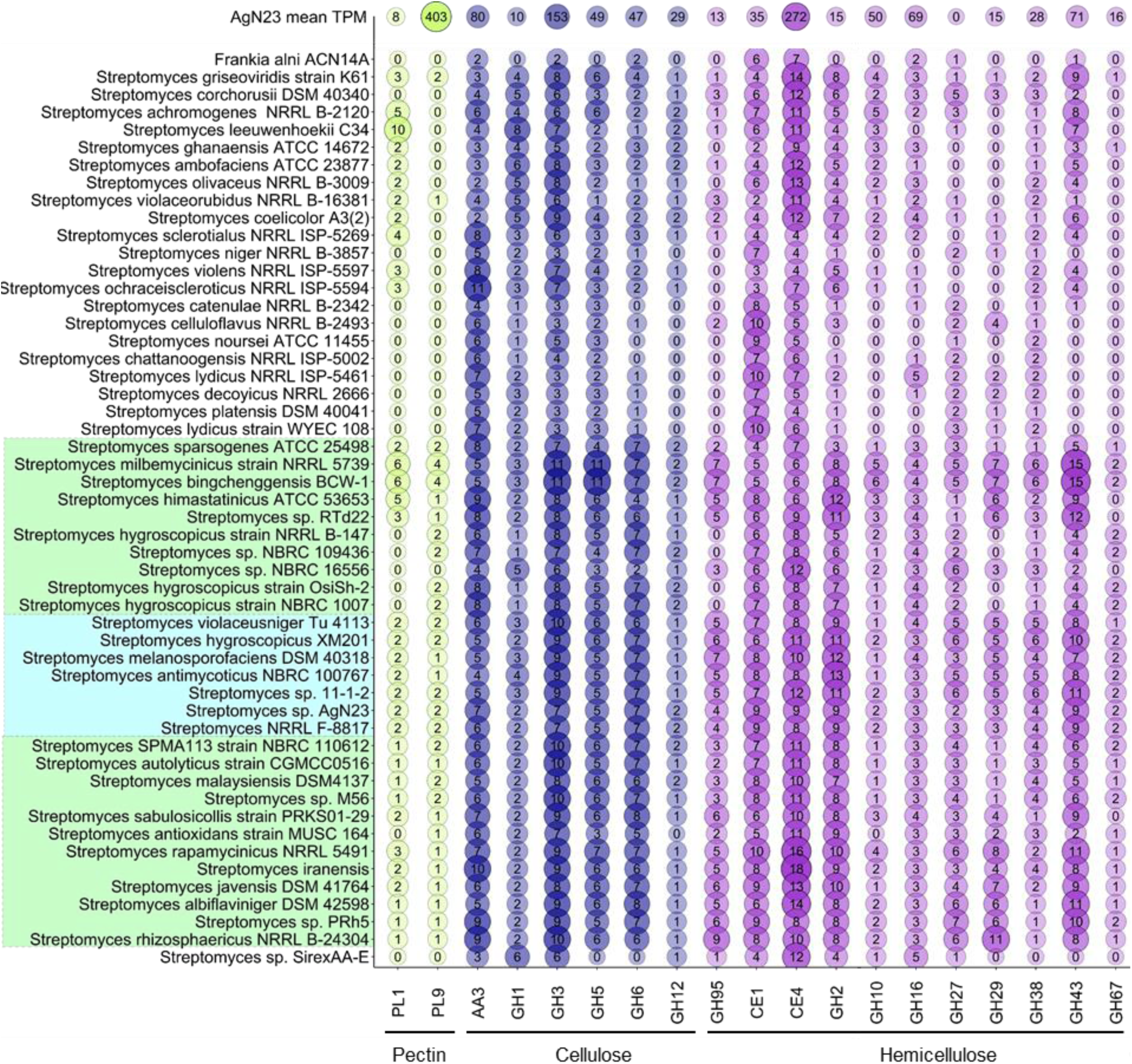
CAZyome associated with degradation of plant cell wall polysaccharides is expanded in AgN23 and related *S. violaceusniger* sp. as compared to outgroup strains. CAZymes families with at least 2 genes in AgN23 are ordered according to their target polymers and the type of enzymatic activity: Glycoside Hydrolases (GHs), Polysaccharide Lyases (PLs), Carbohydrate Esterases (CEs) and Auxiliary Activities (AAs). The transcription of AgN23 CAZyome is displayed as mean Transcript Per Million reads (TPM) for each family (n=3).

We found that AgN23 and other members of the *S. violaceusniger* clade possess two PL9 Polysaccharide Lyases involved in pectin degradation which are absent from 20 of the 22 *Streptomyces* outgroup genomes. Similarly, AgN23 genome harbours 7 GH6 (Glycoside Hydrolases) that are endoglucanases related to cellulose degradation. Such large repertoire of GH6 is frequent in *S. violaceusniger* clade (ANI>85%) whilst 0 to 4 genes were found in the outgroup strains tested. A similar observation could be drawn for GH43 and GH95 involved in hemicellulose catabolism. Strikingly, AgN23 genome encodes 9 GH43 whilst the other *S. violaceusniger* ssp. possess up to 15 of these genes and 8 strains among the outgroup (ANI<85%) had none.

Finally, we analysed the expression pattern of AgN23 CAZyome, consistenly with the whole genome expression analysis, only 3 CAZymes showed no expression in Bennett liquid culture dataset (Supplementary information 5). Strikingly, two genes of the PL9 family (AgN23_3063 and AgN23_1031) which is largely restricted to the *S. violaceusniger* group showed a strong expression at 526 and 279 TPM respectively. These values rank the two genes among the 10% most expressed CAZymes of AgN23. By contrast, the PL1 representative of AgN23_3657 and AgN23_1017 have TPM of 9 and 8 respectively. Regarding the GH6 and GH43 families, we noticed that they are dominated by the expression of one single gene respectively AgN23_4221 (TPM=199) and AgN23_1548 (TPM=537) which also belongs to the top 10% in terms of CAZy expression in AgN23. Taken together, these genomic and transcriptomic data support the hypothesis that AgN23 degrade plant cell wall cellulose, hemicelluloses and pectins to use them as a carbon source, a feature which may represent a major advantage in the rhizosphere niche. This hypothesis prompts the need to investigate CAZyome expression pattern in the frame of plant co-cultivation with AgN23.

### AgN23’s specialised metabolism targets biological functions in plants and fungi

Specialised metabolites are presumed to play key roles in adaption of streptomycetes to their environment. We used antiSMASH 6.0 to detect and annotate AgN23 BGCs according to their similarity to reference clusters deposited in the MIBiG database [24, 25]. Forty-seven BGCs were detected in the AgN23 genome, all of them being expressed during AgN23 cultivation in Bennett liquid medium except for region 47 (Table 2, Supplementary information 6). Only the region 20, putatively involved in pigment biosynthesis stood out in terms of expression with 3330 TPM, all the other ranging between a few to 200 TPM. Twenty BGCs of AgN23 showed at least 50% of similar gene content with MIBiG reference BGCs (Table 2). These candidates BGCs are notably involved in the biosynthesis of volatiles terpenes (regions 3, 10) including geosmin region23), indoles (region 21), ribosomally synthesised and post-translationally modified peptides-like (RiPP-like, regions 5 and 28), ochronotic pigment (region 20) and a bicyclomycin-like antibacterial (region 45). Other BGCs are potentially involved in the production of siderophores including coelichelin and desferrioxamin (region 24, 27, 44) or osmotic and cold stresses protectant (ectoin, region 15). Additional BGCs are likely involved in the regulation of the bacterium life cycle such as spore pigment (region 31), hopene (region 32), and butyrolactone (region 37). BGCs encoded in regions 2, 17, 41 and 42 are similar to the one belonging to the biosynthesis pathways of the antifungal compounds rustmicin, also known as galbonolide A, mediomycin A, nigericin and niphimycins C-E respectively [57-60]. In addition, echoside (region 26), elaiophylin (region 40) and nigericin (region 41) are structural analogues of terfestatin, pteridic acid and monensin respectively, three compounds affecting plant immunity and development [61-65].

**Table 2:**
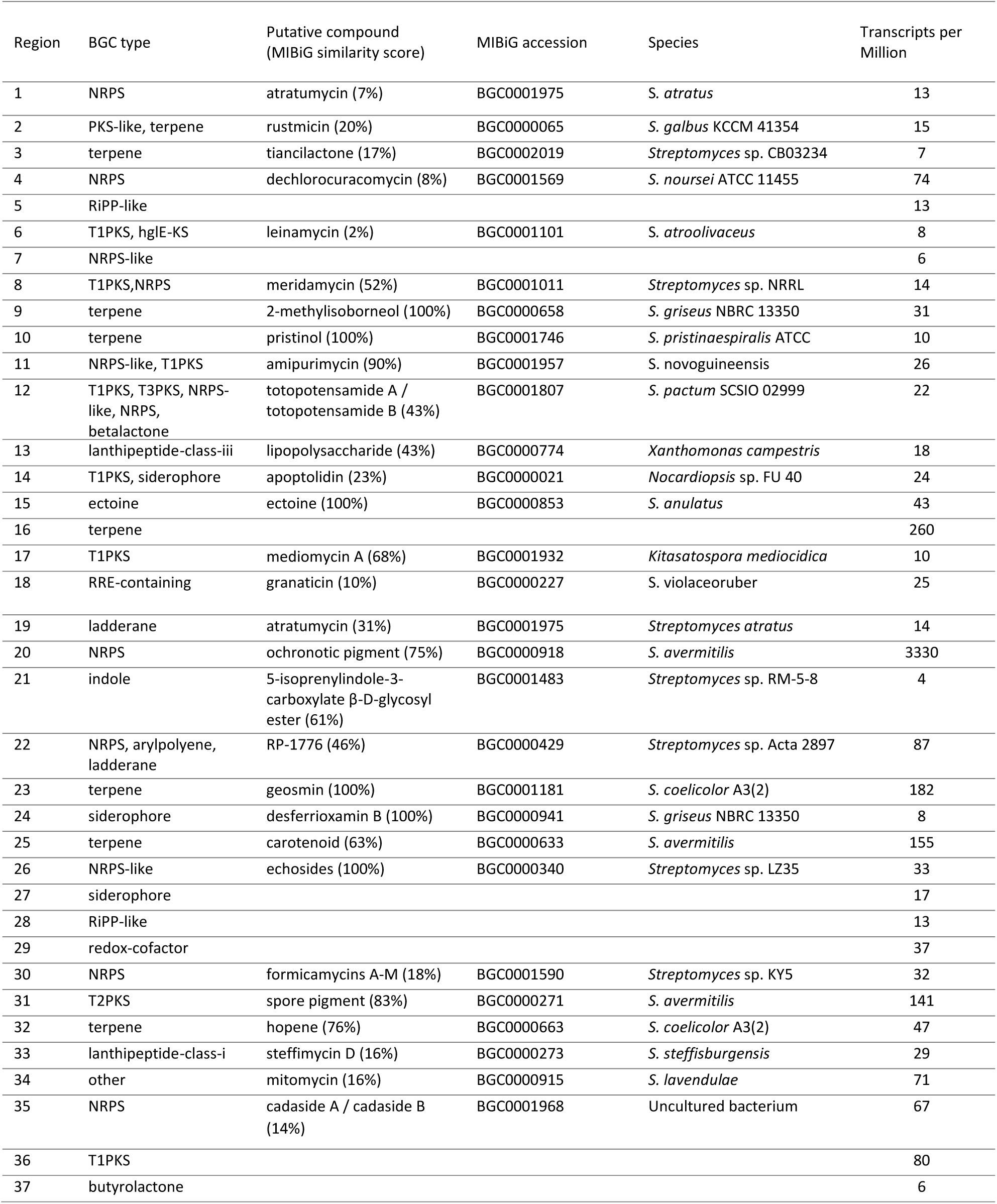
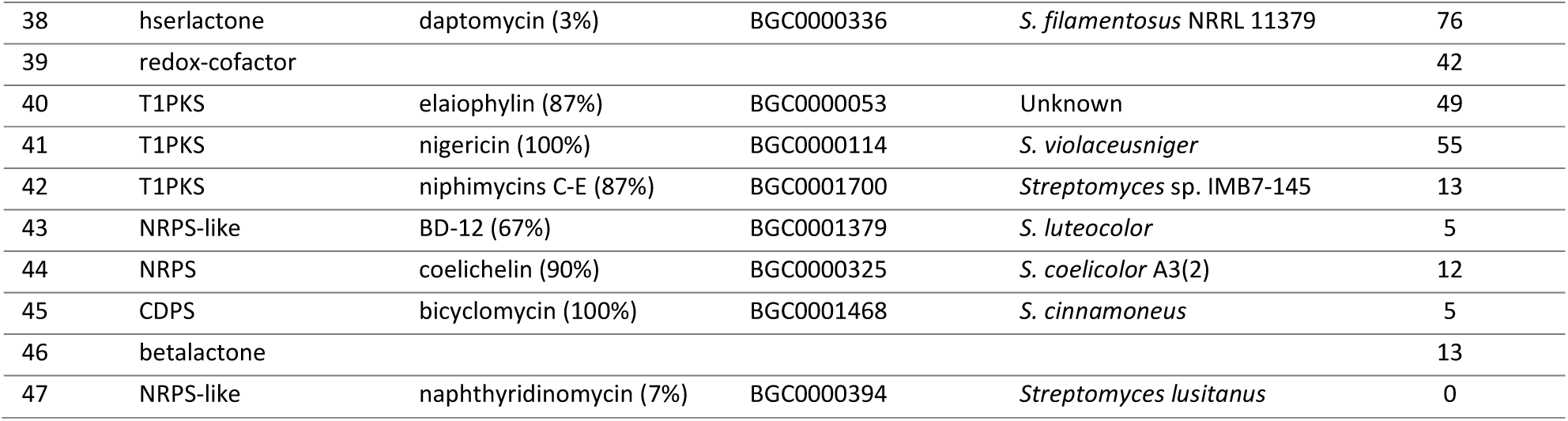
antiSMASH annotation of AgN23 chromosomal regions coding for Biosynthetic Gene Clusters. The functional category of each BGCs was determined by antiSMASH. The BGC type, best hit in the MIBiG database as its percentage of similarity to the query are indicated along with the bacterial strain from whom the cluster was described. The expression level of each BGCs was determined by doing the mean of the expression level in Transcripts Per Million (TPM) of the core biosynthetic genes of each BGC from the RNA-seq data (n=3).

In addition to the BGC-mediated production of specialised metabolites, some *Streptomyces* spp. produce auxin, a phytohormone with a strong impact on root growth [66]. We undertook the annotation of auxin biosynthesis genes of AgN23 by retrieving protein sequences associated to KEGG ontologies leading to the biosynthesis of indole-3 acetic acid (IAA), the most common plant auxin, from its precursor tryptophane through the indole-3-acetamide (IAM) pathway, the indole-3-acetaldoxime/indole-3-acetonitrile (IAN/IAOx) pathway, the tryptamine (TAM) or indole-3-pyruvate (IPyA) pathways both leading to indole-3-acetaldehyde (IAAld) before its conversion to IAA (Supplementary Information 7)[23]. As a result, we identified a complete biosynthetic route for the IAM pathway (AgN23_8393 and AgN23_8392), the TAM-IAAld pathway (AgN23_3181, AgN23_0775, AgN23_0524 and AgN23_0525) and the IPyA to IAAld pathway (AgN23_1600) as well as enzymes converting IAN to IAM (AgN23_1182 and AgN23_1183)(Figure 4). Taken together, these data reveal that AgN23 likely produce specialised metabolites and auxins with a potential to regulate host plant biology along with its root microbiota.

**Figure 4:**
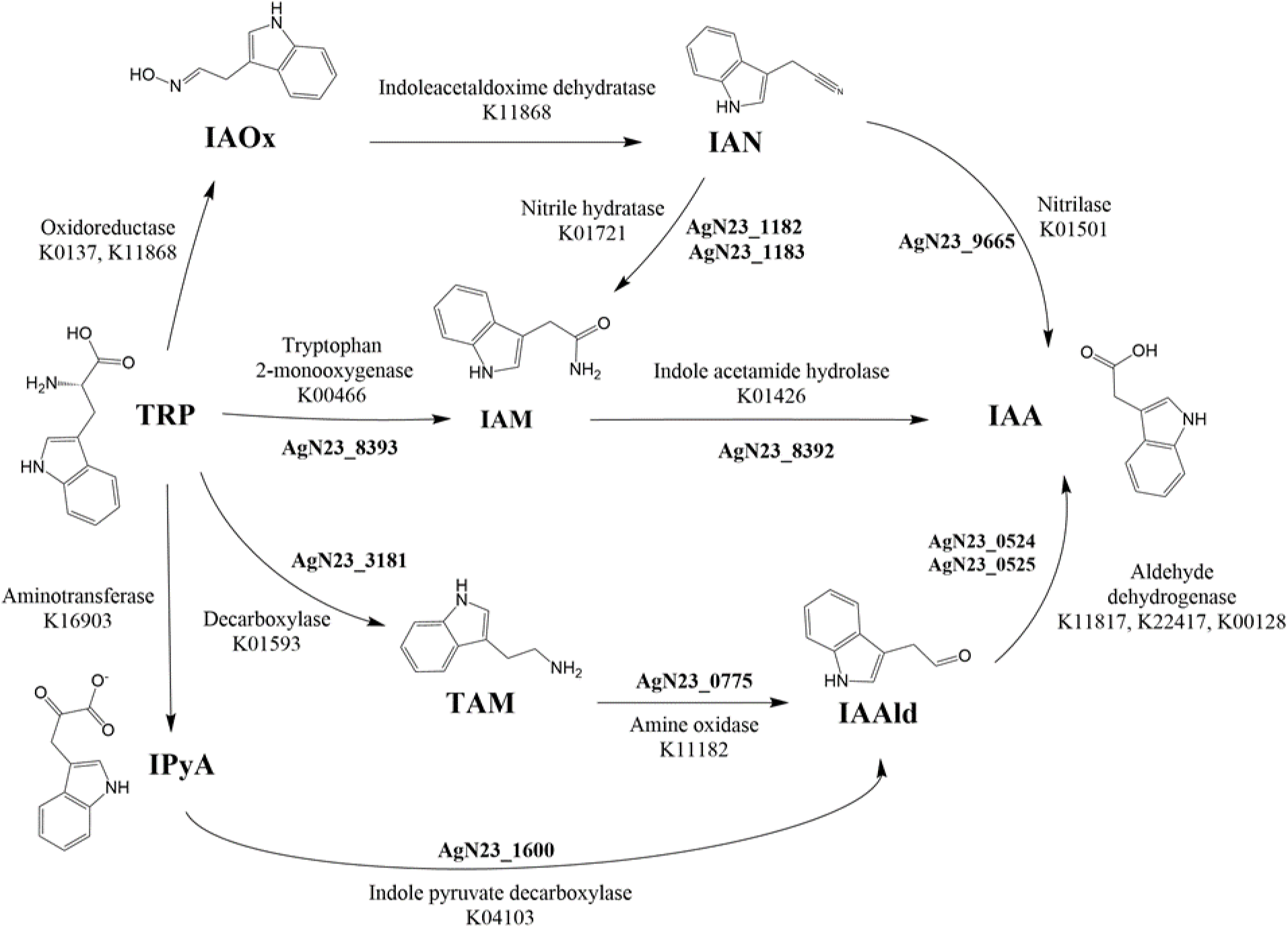
Overview of tryptophan-dependent indole-3 acetic acid (IAA) biosynthesis pathways in AgN23. AgN23 genes encoding putative enzymes and their cognate KEGG annotations are indicated. Compound abbreviations: tryptophan (Trp), indole-3-acetaldoxime (IAOx), indole-3-acetonitrile (IAN), indole-3-acetamide (IAM), indole-3-pyruvate (IPyA), indole-3-acetaldehyde (IAAld) and tryptamine (TAM).

### A core set of BGCs conserved across *S. violaceusniger* species

The *S. violaceusniger* clade hosts several strains possessing BGCs involved in the synthesis of antifungal polyene macrolides such as nigericin, elaiophylin and geldanamycin [32, 56, 67-72]. However, strains sharing phylogenetic vicinity may differ in their specialised metabolism due to variation in their BGCs [73, 74]. To assess the diversity of BGCs within the *S. violaceusniger* clade, we set up a comparative genomic approach based on the chromosome-scale assemblies used for our phylogenetic analysis (Supplementary information 1). antiSMASH retrieved 873 BGC-containing regions in the genomes of AgN23 and the 18 other selected genomes. These sequences were introduced into BiG-SCAPE which grouped them into 415 gene clusters families (GCF) and built a sequence similarity network. To support our BiG-SCAPE analysis, the sequences of the 47 AgN23 BGCs were then compared to the 1,225,071 BGCs of the BiG-FAM database using the BiG-SLICE algorithm [15, 75].

We first observed that BiG-SCAPE distributed the AgN23 BGCs in four categories comprising *S. violaceusniger* specific BGCs (ANI>85%) (Figure 5a), BGCs found in outgroup (ANI<85%) (Figure 5b) and AgN23 BGCs not involved in any GCFs (Figure 5c). In addtion, BiG-SCAPE identified GCFs involving BGCs of others *S. violaceusniger* strains but not from AgN23 (Figure 5d). To back-up the identification of BGCs specific to *S. violaceusniger* (Figure 5a) or unique to AgN23 (Figure 5c), we inspected any potential hits of the corresponding AgN23 BGCs in the BiG-FAM database (Supplementary Information 8). As a result, 6 BGCs found to be *S. violaceusniger* specific in BiG-SCAPE (Figure 5a) actually had outgroup hits in BiG-FAM. Also, one BGC with no hits in BiG-SCAPE had a similar BGC in Big-FAM. The result of BiG-SCAPE and Big-FAM analyses is consolidated in Figure 6.

**Figure 5:**
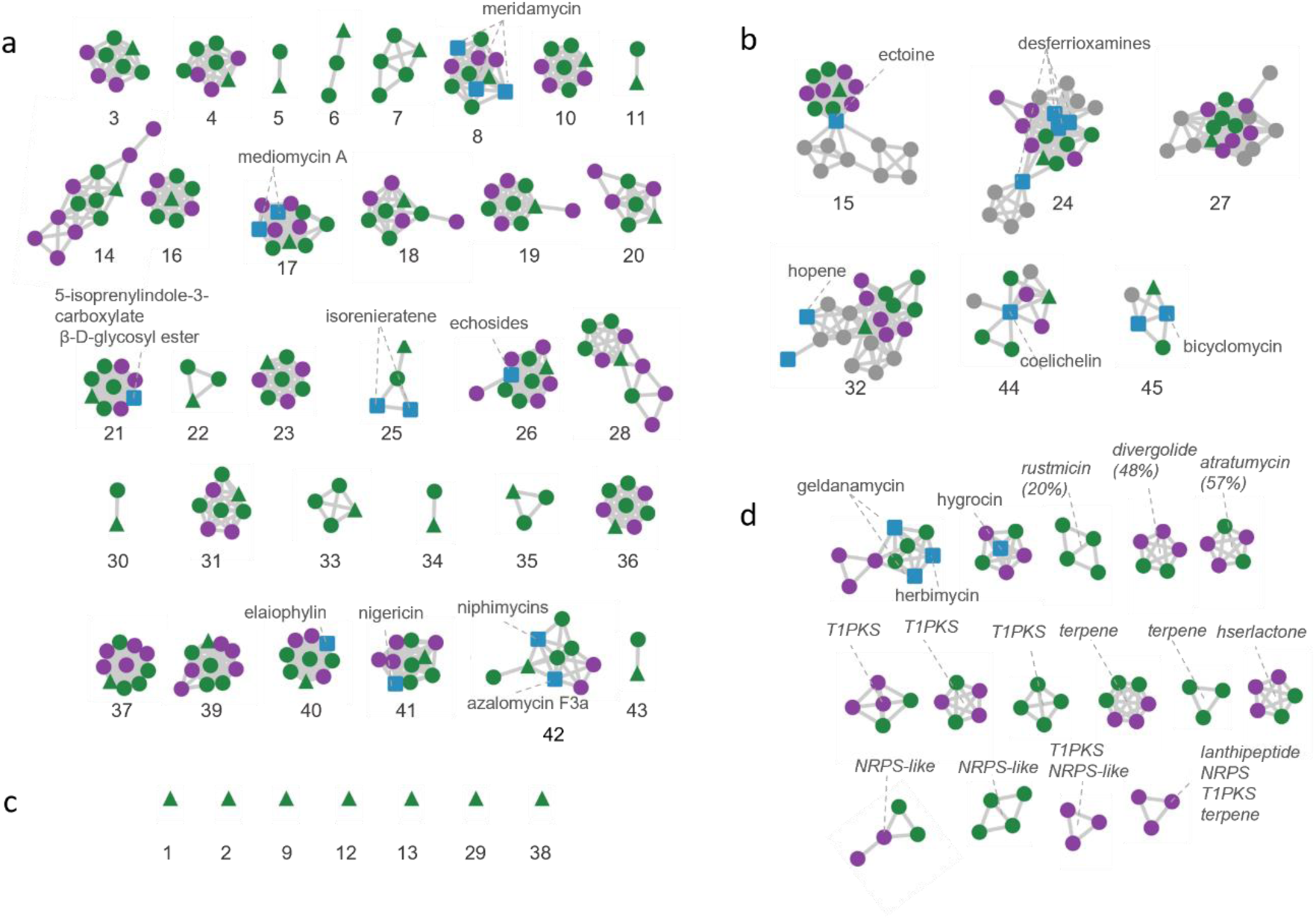
*Streptomyces* sp. AgN23 shares a large set of BGCs specifically found within *S. violaceusniger* calde. A BiG-SCAPE similarity network was built with the BGCs of AgN23 and 18 *Streptomyces* isolates selected both inside and outside *S. violaceusniger* clade. Each node represents a BGC-containing region from one genome and clusters of nodes are based on BGCs similarity. AgN23 BGCs are displayed as green triangles. Blue squares represent curated sequences from the MiBIG database. The round node corresponds to BGCs from the 18 selected strains. These nodes are green from strains with ANI>95% with AgN23, strains with ANI>85% ae in purple and and strains with ANI<85% are in grey. a) Clusters of BGCs specific to AgN23 and strains with ANI>85%. b) Clusters of BGCs including AgN23 and strains with ANI<85%. c) AgN23 BGCs which were not assigned to any cluster. d) Clusters of BGCs from strains with ANI>85% with AgN23 for whom no homologous BGCs were found in AgN23.

**Figure 6:**
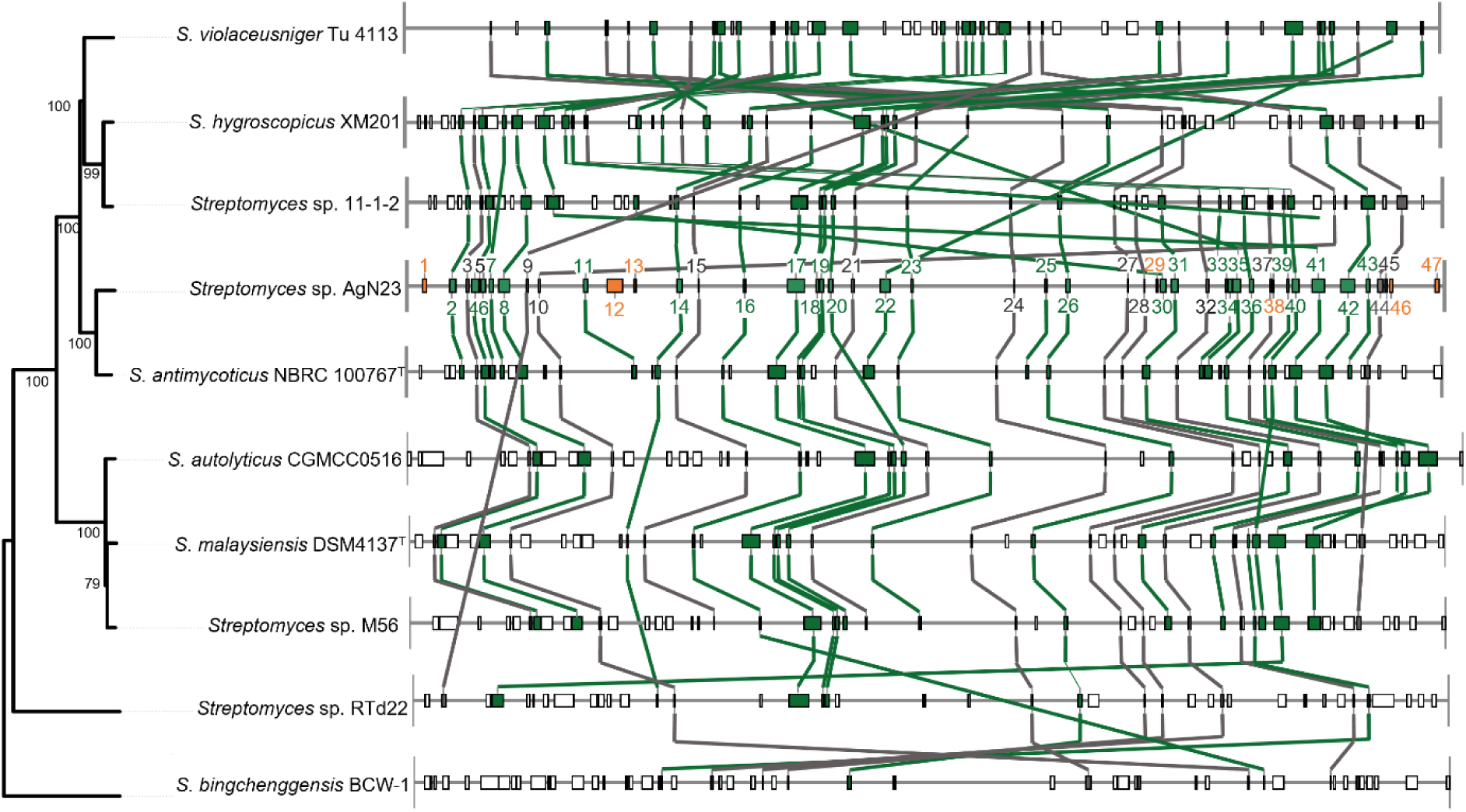
Synteny analysis of the BGC-containing regions in genomes of *S. violaceusniger* clade. Each block represent a BGC-containing region on a chromosme. The BGCs are linked in green between different genomes if they were clustered by BiG-SCAPE and confirmed to be specific to *S. violaceusniger* by BiG-FAM analysis. BGCs linked in grey were clustered by BiG-SCAPE and found outside the *S. violaceusniger* clade. Orange blocks of AgN23 represent BGCs unique to the strain. Tree inferred from Genome BLAST Distance Phylogeny (GBDP) distances calculated from fully-assembled chromosome of isolates from the *S. violaceuniger* clade. The numbers above branches are GBDP pseudo-bootstrap support values > 60 % from 100 replications.

These analyses revealed that BGCs involved in the biosynthesis of geldanamycin, a hallmark phytotoxic, antifungal and antibacterial compound of *S. violaceusniger* strains was detected in 6 strains but neither in AgN23 nor its closest neighbour *S. antimycoticus* (Figure 5d)[76]. Also, BiG-SCAPE analysis clustered regions containing rustmicin-like BGCs from four *S. violaceusniger* strains but none of both tools did not connect the region 2 of AgN23 to it. Though, that region contains a BGC annotated as rustmicin by antiSMASH (Table 2). Since the organisation of the five BGCs are very similar to the subcluster GalA-E of *S. galbus* responsible for the biosynthesis of the macrolactone ring of the compound, we propose that the biosynthesis of a rustmicin-like compound is commonly found within the *S. violaceusniger* clade [77](Figure 7). Importantly both BiG-SCAPE and BiG-FAM confirmed the conservation of BGCs involved in antifungal activity (regions 2, 17, 41 and 42) and plant bioactive metabolites (regions 26, 40 and 41) which we identified in AgN23 and the *S. violaceusniger* clade. This suggests that this specialisation could have arisen during the *Streptomyces* genus radiation and may result, or have participated, in the *S. violaceusniger* clade adapatation to the plant rhizosphere [78-80].

**Figure 7:**
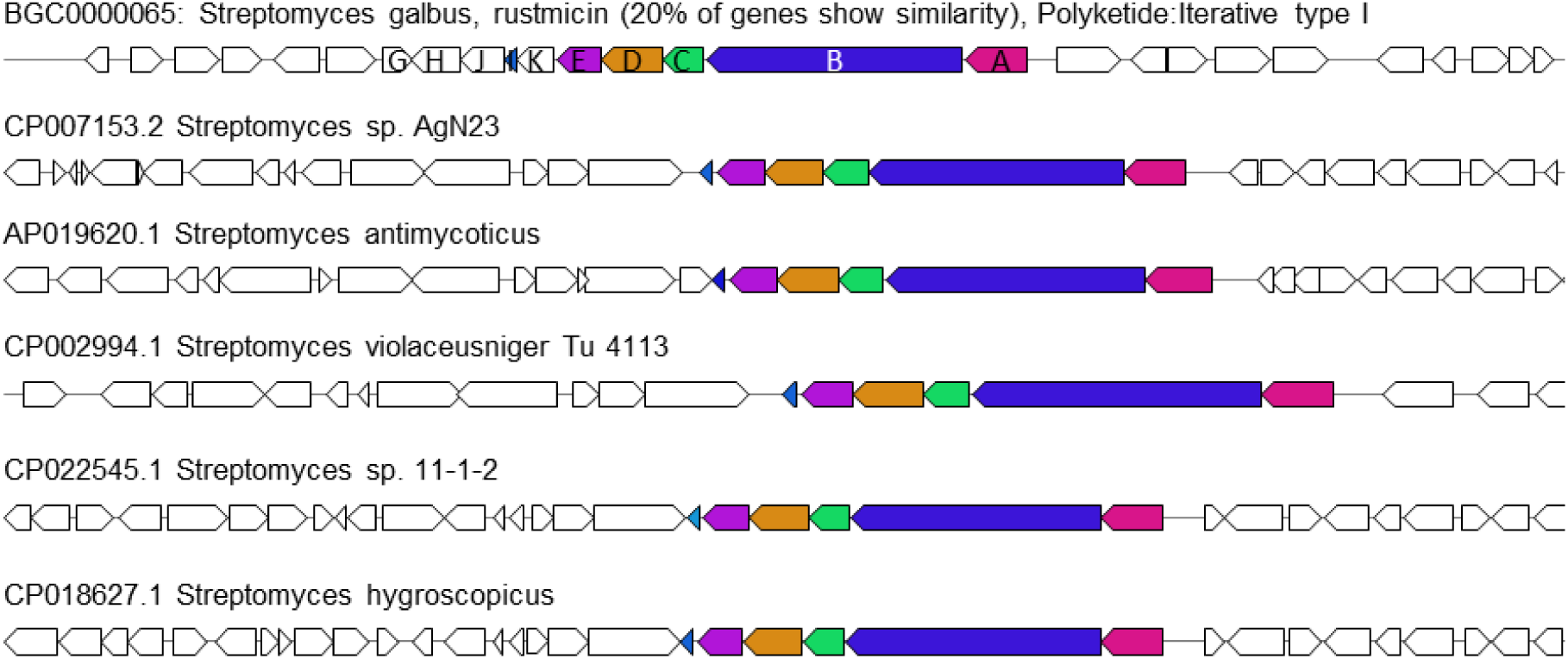
rustmicin/galbonolide ABCDE biosynthetic gene cluster is conserved across *S. violaceusniger* clade. Comparison of the rustmicin BGC sequence from MIBiG (BGC0000065) with genomic region 2 of AgN23 and corresponding regions of four others *S. violaceusniger* genomes. Coloured genes are putative homologs according to antiSMASH with ClusterBLAST *e* -value < 1E-05, an identity > 30% and a query cover > 25%.

### BGCs synteny and horizontal transfers accross *S. violaceusniger* clade chromosomes

*Streptomyces* chromosome organisation consists in a central conserved genome whilst terminal sequences contain more variable gene content described as accessory genome [81-83]. This accessory genome undergoes frequent rearrangement, amplification and deletion events as well as interspecies homologous recombination [79, 84-89]. We decided to investigate commonalities and differences in the BGC organisation accross *S. violaceusniger* clade and to pinpoint original features of our newly sequenced AgN23 strain.

We found that the core set of 26 BGCs found in *S. violaceusnige*r clade is syntenically positionned in « left » and « right » arms of the chromosome or in the central region in AgN23, *Streptomyces* sp. 11-1-2, *S. antimycoticus* NBRC10767, *S. autolyticus* CGMCC0516, *S. malaysiensis* DSM4137 and *Streptomyces* sp. M56 (Figure 6). By contrast, discrepancies in BGCs positions exist between this latter group of genomes and *S. violaceusniger* Tü 4113 as well as *S. hygroscopicus* XM201 suggesting these strains underwent drastic chromosome rearrangements but still retained BGCs connected to the synthesis of plant and fungal bioactive metabolites.

The regions 1, 46 and 47 which harbour original BGCs according to antiSMASH (<10% similarity with MIBiG) and the BiG-SCAPE analysis are located on the extremities left and right arms of the AgN23 chromosome and syntenic strains. The region 45, contains a BGC with 100% similarity to the biosynthetic pathway of the antimicrobial bicyclomycin, which is absent from other *S. violaceusniger* strains but is found outside of the clade (Table 2, Supplementary information 8). Thus, regions 45 may have been acquired by AgN23 through horizontal gene transfer. In conclusion, our synteny analysis is consistent with a preferential accumulation of unique or horizontally transferred BGCs on chromosome extremities of AgN23. Conversely the biosynthesis of specialised metabolites such as niphimycin or galbonolide was reported outside the *S. violaceusniger* clade, suggesting that some of these specific BGCs may have been horizontally transferred outside of the clade [57, 60]. Taken together, this data supports the view that AgN23 accessory BGCs are concentrated on chromosome extremities.

## Conclusions

Plant roots recruit abundant and diverse consortia of microorganisms whilst exploring the soil. The relevance of the root microbiota in plant nutrition and resistance to stresses is currently unveiled through metabarcoding studies of plant microbiota. *Streptomyces* genus constitutes one of the most prominent bacterial genera colonising plant roots [7]. However, such studies do not inform on the species diversity of *Streptomyces* and the genomic features enabling their colonisation of plant roots. Therefore, genome sequencing of root associated *Streptomyces* is an important step toward the description of their molecular interaction with host plant. Here we produced a complete chromosome sequence of *Streptomyces* sp. AgN23, a strain isolated from grapevine rhizosphere and shown to elicit plant defence responses [16]. A multi-locus sequence typing phylogeny based on AgN23 genome sequence showed that it belongs to the *S. violaceusniger* clade from whom several other strains have been isolated from the rhizosphere of diverse plants around the world. We found that AgN23 genome, as well as other representative strains of the *S. violaceusniger* clade is rich in CAZymes able to degrade plant-derived carbohydrates and in BGCs producing specialised metabolite potentially involved in the interaction with the plant. In addition, this phylogenetic lineage is considered as having a high potential in terms of specialised metabolism as its members have a large genome between 10.7 and 12.7 Mb and possess from 45 to 55 BGCs [90]. The wealth of genomic data available in this clade allowed us to unveil common trends in the BGCs specifically found in *S. violaceusniger* isolates. The ability of these strains to produce antifungal compounds (nigericin, niphimycin, mediomycin, rustmicin) as well as elicitors of plant defence (nigericin) strongly suggests that they play an important role in protection from pathogens. In addition, echosides and elaiophylin from whom structural analogues have been shown to interfere with auxin like responses suggest they may interfere with plant development [62, 64]. Thorough comparisons in the BGC content and the chromosome organisation of AgN23 with other *S. violaceusniger* sp. also enabled to pinpoint BGCs gains (i.e. bicyclomycin) and losses (i.e. geldanamycin) which may account for specific activities of the strain within the plant root microbiota.

## Author statements

### Authors and contributors

D. Gayrard performed the research and wrote the manuscript. M. Veyssière, C. Nicolle, K. Adam, Y. Martinez, C. Vandecasteele and M. Vidal performed the research. B. Dumas and T. Rey designed the research and wrote the manuscript.

### Conflicts of interest

The following information may be foreseen as competing interest. B. Dumas is one of inventors of the patent WO2015044585A1 relating the use of AgN23 in Agriculture. T. Rey and D. Gayrard are full-time researchers in the AgChem company De Sangosse (Pont-Du-Casse, France), which registers and markets crop-protection products.

### Funding information

This work was funded by the Fond Unique Interministériels (NEOPROTEC project), the Fonds Européen de Développement Économique et Régional (FEDER), the Agence Nationale de la Recherche (LabCom BioPlantProtec ANR-14-LAB7-0001 and STREPTOCONTROL ANR-17-CE20-0030) and the Région Occitanie (projet GRAINE-BioPlantProducts). The Laboratoire de Recherche en Sciences Végétales (LRSV) belongs to the TULIP Laboratoire d’Excellence (ANR-10-LABX-41). Work performed in the GeT core facility, Toulouse, France (https://get.genotoul.fr) was supported by the France Génomique National infrastructure, funded as part of the “Investissement d’Avenir” program managed by the Agence Nationale de la Recherche (contract ANR-10-INBS-09) and by the GET-PACBIO program (FEDER Programme opérationnel FEDER-FSE MIDI-PYRENEES ET GARONNE 2014-2020). D. Gayrard was funded by the Agence Nationale de la Recherche Technique, with the Convention Industrielle de Formation par la Recherche and Association Nationale de la Recherche et de la Technologie (grant number 2016/1297).

## Acknowledgements

We warmly thanks Olivier Bouchez for helpful discussions on Next Generation Sequencing strategy throughout the project. We are grateful to Sylvie Lautru and Jean-Luc Pernodet (I2BC, Paris) for helpful discussions. We thank Thierry Grollier and Valérie Arnal for helpful comments on earlier version of the manuscript.

## Notes

### Summary of Updates

Mistakes in the material and methods section have been corrected. The figures have been modified with expanded datasets.

## References

1. Vallenet D, Calteau A, Dubois M, Amours P, Bazin A et al. MicroScope: an integrated platform for the annotation and exploration of microbial gene functions through genomic, pangenomic and metabolic comparative analysis. Nucleic Acids Res 2020;48(D1):D579–D589.

2. Bush MJ, Tschowri N, Schlimpert S, Flärdh K, Buttner MJ. c-di-GMP signalling and the regulation of developmental transitions in streptomycetes. Nat Rev Microbiol 2015;13(12):749–760.

3. Book AJ, Lewin GR, Mcdonald BR, Takasuka TE, Wendt-Pienkowski E et al. Evolution of High Cellulolytic Activity in Symbiotic Streptomyces through Selection of Expanded Gene Content and Coordinated Gene Expression. PLOS Biology 2016;14(6):e1002475.

4. Moumbock AFA, Gao M, Qaseem A, Li J, Kirchner PA et al. StreptomeDB 3.0: an updated compendium of streptomycetes natural products. Nucleic Acids Res 2021;49(D1):D600–D604.

5. Barka EA, Vatsa P, Sanchez L, Gaveau-Vaillant N, Jacquard C et al. Taxonomy, Physiology, and Natural Products of Actinobacteria. Microbiol Mol Biol Rev 2016;80(1):1–43.

6. Lee N, Kim W, Hwang S, Lee Y, Cho S et al. Thirty complete Streptomyces genome sequences for mining novel secondary metabolite biosynthetic gene clusters. Sci Data 2020;7(1):55.

7. Lundberg DS, Lebeis SL, Paredes SH, Yourstone S, Gehring J et al. Defining the core Arabidopsis thaliana root microbiome. Nature 2012;488(7409):86–90.

8. Fitzpatrick CR, Copeland J, Wang PW, Guttman DS, Kotanen PM et al. Assembly and ecological function of the root microbiome across angiosperm plant species. Proc Natl Acad Sci U S A 2018;115(6):E1157–E1165.

9. Vurukonda SSKP, Giovanardi D, Stefani E. Plant Growth Promoting and Biocontrol Activity of Streptomyces spp. as Endophytes. Int J Mol Sci 2018;19(4).

10. Hamedi J, Mohammadipanah F. Biotechnological application and taxonomical distribution of plant growth promoting actinobacteria. Journal of industrial microbiology & biotechnology 2015;42(2):157–171.

11. Rey T, Dumas B. Plenty Is No Plague: Streptomyces Symbiosis with Crops. Trends Plant Sci 2017;22(1):30–37.

12. Viaene T, Langendries S, Beirinckx S, Maes M, Goormachtig S. Streptomyces as a plant’s best friend? FEMS Microbiol Ecol 2016;92(8).

13. Medema MH. The year 2020 in natural product bioinformatics: an overview of the latest tools and databases. Nat Prod Rep 2021;38(2):301–306.

14. Tracanna V, de Jong A, Medema MH, Kuipers OP. Mining prokaryotes for antimicrobial compounds: from diversity to function. FEMS Microbiol Rev 2017;41(3):417–429.

15. Kautsar SA, Blin K, Shaw S, Weber T, Medema MH. BiG-FAM: the biosynthetic gene cluster families database. Nucleic Acids Res 2021;49(D1):D490–D497.

16. Vergnes S, Gayrard D, Veyssiere M, Toulotte J, Martinez Y et al. Phyllosphere colonisation by a soil Streptomyces sp. promotes plant defense responses against fungal infection. Mol Plant Microbe Interact 2019; 33(2):223–234.

17. Errakhi R, Bouteau F, Lebrihi A, Barakate M. Evidences of biological control capacities of Streptomyces spp. against Sclerotium rolfsii responsible for damping-off disease in sugar beet (Beta vulgaris L.). World Journal of Microbiology and Biotechnology 2007;23(11):1503–1509.

18. Shirling EB, Gottlieb D. Methods for characterization of Streptomyces species. International Journal of Systematic Bacteriology 1966. p. 313–340.

19. Kieser T, Bibb MJ, Buttner MJ, Chater KF, Hopwood DA. Practical streptomyces genetics: John Innes Foundation Norwich; 2000.

20. Parks DH, Imelfort M, Skennerton CT, Hugenholtz P, Tyson GW. CheckM: assessing the quality of microbial genomes recovered from isolates, single cells, and metagenomes. Genome Research 2015;25(7):1043–1055.

21. Simão FA, Waterhouse RM, Ioannidis P, Kriventseva EV, Zdobnov EM. BUSCO: assessing genome assembly and annotation completeness with single-copy orthologs. Bioinformatics 2015;31(19):3210–3212.

22. Yin Y, Mao X, Yang J, Chen X, Mao F et al. dbCAN: a web resource for automated carbohydrate-active enzyme annotation. Nucleic Acids Res 2012;40(W1):W445–W451.

23. Zhang P, Jin T, Kumar Sahu S, Xu J, Shi Q et al. The Distribution of Tryptophan-Dependent Indole-3-Acetic Acid Synthesis Pathways in Bacteria Unraveled by Large-Scale Genomic Analysis. Molecules 2019;24(7).

24. Blin K, Shaw S, Steinke K, Villebro R, Ziemert N et al. antiSMASH 5.0: updates to the secondary metabolite genome mining pipeline. Nucleic Acids Res 2019;47(W1):W81–W87.

25. Epstein SC, Charkoudian LK, Medema MH. A standardized workflow for submitting data to the Minimum Information about a Biosynthetic Gene cluster (MIBiG) repository: prospects for research-based educational experiences. Stand Genomic Sci 2018;13:16.

26. Navarro-Muñoz JC, Selem-Mojica N, Mullowney MW, Kautsar SA, Tryon JH et al. A computational framework to explore large-scale biosynthetic diversity. Nat Chem Biol 2020;16(1):60–68.

27. Kautsar SA, Blin K, Shaw S, Weber T, Medema MH. BiG-FAM: the biosynthetic gene cluster families database. Nucleic Acids Res 2021b;49:D490–D497.

28. Klassen JL, Lee SR, Poulsen M, Beemelmanns C, Kim KH. Efomycins K and L From a Termite-Associated. Front Microbiol 2019;10:1739.

29. Díaz-Cruz GA, Liu J, Tahlan K, Bignell DRD. Nigericin and Geldanamycin Are Phytotoxic Specialized Metabolites Produced by the Plant Pathogen. Microbiol Spectr 2022:e0231421.

30. Kusuma AB, Nouioui I, Goodfellow M. Genome-based classification of the Streptomyces violaceusniger clade and description of Streptomyces sabulosicollis sp. nov. from an Indonesian sand dune. Antonie Van Leeuwenhoek 2021;114(6):859–873.

31. Kumar Y, Aiemsum-Ang P, Ward AC, Goodfellow M. Diversity and geographical distribution of members of the Streptomyces violaceusniger 16S rRNA gene clade detected by clade-specific PCR primers. FEMS Microbiol Ecol 2007;62(1):54–63.

32. Goodfellow M, Kumar Y, Labeda DP, Sembiring L. The Streptomyces violaceusniger clade: a home for Streptomycetes with rugose ornamented spores. Antonie Van Leeuwenhoek 2007;92(2):173–199.

33. Alanjary M, Steinke K, Ziemert N. AutoMLST: an automated web server for generating multi-locus species trees highlighting natural product potential. Nucleic Acids Res 2019;47(W1):W276–W282.

34. Lefort V, Desper R, Gascuel O. FastME 2.0: A Comprehensive, Accurate, and Fast Distance-Based Phylogeny Inference Program. Mol Biol Evol 2015;32(10):2798–2800.

35. Meier-Kolthoff JP, Göker M. TYGS is an automated high-throughput platform for state-of-the-art genome-based taxonomy. Nat Commun 2019;10(1):2182.

36. Li H. Aligning sequence reads, clone sequences and assembly contigs with BWA-MEM. arXiv preprint 13033997 2013.

37. Li H, Handsaker B, Wysoker A, Fennell T, Ruan J et al. The sequence alignment/map format and SAMtools. Bioinformatics 2009;25(16):2078–2079.

38. Quinlan AR, Hall IM. BEDTools: a flexible suite of utilities for comparing genomic features. Bioinformatics 2010;26(6):841–842.

39. Labeda DP, Goodfellow M, Brown R, Ward AC, Lanoot B et al. Phylogenetic study of the species within the family Streptomycetaceae. Antonie Van Leeuwenhoek 2012;101(1):73–104.

40. Jain C, Rodriguez-R LM, Phillippy AM, Konstantinidis KT, Aluru S. High throughput ANI analysis of 90K prokaryotic genomes reveals clear species boundaries. Nat Commun 2018;9(1):5114.

41. Komaki H, Tamura T. Reclassification of Streptomyces castelarensis and Streptomyces sporoclivatus as later heterotypic synonyms of Streptomyces antimycoticus. International Journal of Systematic and Evolutionary Microbiology 2020.

42. Caicedo-Montoya C, Manzo-Ruiz M, Ríos-Estepa R. Pan-Genome of the Genus. Front Microbiol 2021;12:677558.

43. Goodfellow M, Kumar Y, Labeda DP, Sembiring L. The Streptomyces violaceusniger clade: a home for streptomycetes with rugose ornamented spores. Antonie van Leeuwenhoek 2007;92(2):173–199.

44. Yin M, Jiang M, Ren Z, Dong Y, Lu T. The complete genome sequence of Streptomyces autolyticus CGMCC 0516, the producer of geldanamycin, autolytimycin, reblastatin and elaiophylin. J Biotechnol 2017;252:27–31.

45. Ser H-L, Tan LT-H, Palanisamy UD, Abd Malek SN, Yin W-F et al. Streptomyces antioxidans sp. nov., a Novel Mangrove Soil Actinobacterium with Antioxidative and Neuroprotective Potentials. Frontiers in Microbiology 2016;7.

46. Hamedi J, Mohammadipanah F, Klenk HP, Potter G, Schumann P et al. Streptomyces iranensis sp. nov., isolated from soil. Int J Syst Evol Microbiol 2010;60(Pt 7):1504–1509.

47. Chagas FO, Ruzzini AC, Bacha LV, Samborskyy M, Conti R et al. Genome Sequence of Streptomyces sp. Strain RTd22, an Endophyte of the Mexican Sunflower. Genome Announc 2016;4(4).

48. Yang H, Zhang Z, Yan R, Wang Y, Zhu D. Draft Genome Sequence of Streptomyces sp. Strain PRh5, a Novel Endophytic Actinomycete Isolated from Dongxiang Wild Rice Root. Genome Announc 2014;2(2).

49. Bown L, Bignell DRD. Draft Genome Sequence of the Plant Pathogen. Genome Announc 2017;5(37).

50. Zeng J, Xu T, Cao L, Tong C, Zhang X et al. The Role of Iron Competition in the Antagonistic Action of the Rice Endophyte Streptomyces sporocinereus OsiSh-2 Against the Pathogen Magnaporthe oryzae. Microb Ecol 2018;76(4):1021–1029.

51. Xu T, Li Y, Zeng X, Yang X, Yang Y et al. Isolation and evaluation of endophytic Streptomyces endus OsiSh-2 with potential application for biocontrol of rice blast disease. J Sci Food Agric 2017;97(4):1149–1157.

52. Cao L, Gao Y, Yu J, Niu S, Zeng J et al. Streptomyces hygroscopicus OsiSh-2-induced mitigation of Fe deficiency in rice plants. Plant Physiol Biochem 2021;158:275–283.

53. Gao Y, Ning Q, Yang Y, Liu Y, Niu S et al. Endophytic. mBio 2021:e0156621.

54. Xu T, Cao L, Zeng J, Franco CMM, Yang Y et al. The antifungal action mode of the rice endophyte Streptomyces hygroscopicus OsiSh-2 as a potential biocontrol agent against the rice blast pathogen. Pestic Biochem Physiol 2019;160:58–69.

55. Komaki H, Ichikawa N, Oguchi A, Hamada M, Harunari E et al. Draft genome sequence of Streptomyces sp. TP-A0867, an alchivemycin producer. Stand Genomic Sci 2016;11:85.

56. Kumar Y, Goodfellow M. Five new members of the Streptomyces violaceusniger 16S rRNA gene clade: Streptomyces castelarensis sp. nov., comb. nov., Streptomyces himastatinicus sp. nov., Streptomyces mordarskii sp. nov., Streptomyces rapamycinicus sp. nov. and Streptomyces ruanii sp. nov. Int J Syst Evol Microbiol 2008;58(Pt 6):1369–1378.

57. Hu Y, Wang M, Wu C, Tan Y, Li J et al. Identification and Proposed Relative and Absolute Configurations of Niphimycins C-E from the Marine-Derived Streptomyces sp. IMB7-145 by Genomic Analysis. J Nat Prod 2018;81(1):178–187.

58. Harvey BM, Mironenko T, Sun Y, Hong H, Deng Z et al. Insights into polyether biosynthesis from analysis of the nigericin biosynthetic gene cluster in Streptomyces sp. DSM4137. Chem Biol 2007;14(6):703–714.

59. Cai P, Kong F, Fink P, Ruppen ME, Williamson RT et al. Polyene Antibiotics from Streptomyces mediocidicus. Journal of Natural Products 2007;70(2):215–219.

60. Kim HJ, Karki S, Kwon SY, Park SH, Nahm BH et al. A single module type I polyketide synthase directs de novo macrolactone biogenesis during galbonolide biosynthesis in Streptomyces galbus. J Biol Chem 2014;289(50):34557–34568.

61. Hayashi K, Yamazoe A, Ishibashi Y, Kusaka N, Oono Y et al. Active core structure of terfestatin A, a new specific inhibitor of auxin signaling. Bioorg Med Chem 2008;16(9):5331–5344.

62. Wang X, Reynolds AR, Elshahawi SI, Shaaban KA, Ponomareva LV et al. Terfestatins B and C, New p-Terphenyl Glycosides Produced by Streptomyces sp. RM-5-8. Org Lett 2015;17(11):2796–2799.

63. Yamazoe A, Hayashi K, Kepinski S, Leyser O, Nozaki H. Characterization of terfestatin A, a new specific inhibitor for auxin signaling. Plant Physiol 2005;139(2):779–789.

64. Igarashi Y, Iida T, Yoshida R, Furumai T. Pteridic acids A and B, novel plant growth promoters with auxin-like activity from Streptomyces hygroscopicus TP-A0451. J Antibiot (Tokyo) 2002;55(8):764–767.

65. Maintz J, Suliman M, Joglekar S, Halder V, Kombrink E et al. Chemical Activation of EDS1/PAD4 Signaling Leading to Pathogen Resistance in Arabidopsis. Plant and Cell Physiology 2018;59(8):1592–1607.

66. Rashad FM, Fathy HM, El-Zayat AS, Elghonaimy AM. Isolation and characterization of multifunctional Streptomyces species with antimicrobial, nematicidal and phytohormone activities from marine environments in Egypt. Microbiol Res 2015;175:34–47.

67. Nagpure A, Choudhary B, Gupta RK. Mycolytic enzymes produced by Streptomyces violaceusniger and their role in antagonism towards wood-rotting fungi. J Basic Microbiol 2014;54(5):397–407.

68. Riclea R, Citron CA, Rinkel J, Dickschat JS. Identification of isoafricanol and its terpene cyclase in Streptomyces violaceusniger using CLSA-NMR. Chem Commun (Camb) 2014;50(32):4228–4230.

69. Chen X, Zhang B, Zhang W, Wu X, Zhang M et al. Genome Sequence of Streptomyces violaceusniger Strain SPC6, a Halotolerant Streptomycete That Exhibits Rapid Growth and Development. Genome Announc 2013;1(4).

70. Kang MJ, Strap JL, Crawford DL. Isolation and characterization of potent antifungal strains of the Streptomyces violaceusniger clade active against Candida albicans. J Ind Microbiol Biotechnol 2010;37(1):35–41.

71. Hayakawa M, Yoshida Y, Iimura Y. Selective isolation of bioactive soil actinomycetes belonging to the Streptomyces violaceusniger phenotypic cluster. J Appl Microbiol 2004;96(5):973–981.

72. Hayashi K, Hashimoto M, Shigematsu N, Nishikawa M, Ezaki M et al. WS9326A, a novel tachykinin antagonist isolated from Streptomyces violaceusniger no. 9326. I. Taxonomy, fermentation, isolation, physico-chemical properties and biological activities. J Antibiot (Tokyo) 1992;45(7):1055–1063.

73. Martinet L, Naômé A, Baiwir D, De Pauw E, Mazzucchelli G et al. On the Risks of Phylogeny-Based Strain Prioritization for Drug Discovery:. Biomolecules 2020;10(7).

74. Choudoir MJ, Pepe-Ranney C, Buckley DH. Diversification of Secondary Metabolite Biosynthetic Gene Clusters Coincides with Lineage Divergence in Streptomyces. Antibiotics (Basel) 2018;7(1).

75. Kautsar SA, van der Hooft JJJ, de Ridder D, Medema MH. BiG-SLiCE: A highly scalable tool maps the diversity of 1.2 million biosynthetic gene clusters. Gigascience 2021;10(1).

76. Heisey RM, Putnam AR. Herbicidal effects of geldanamycin and nigericin, antibiotics from Streptomyces hygroscopicus. J Nat Prod 1986;49(5):859–865.

77. Kim H-JJ, Karki S, Kwon S-YY, Park S-HH, Nahm B-HH et al. A single module type I polyketide synthase directs de Novo macrolactone biogenesis during galbonolide biosynthesis in Streptomyces galbus. Journal of Biological Chemistry 2014;289(50):34557–34568.

78. Doroghazi JR, Buckley DH. Intraspecies comparison of Streptomyces pratensis genomes reveals high levels of recombination and gene conservation between strains of disparate geographic origin. BMC Genomics 2014;15:970.

79. Doroghazi JR, Buckley DH. Widespread homologous recombination within and between Streptomyces species. ISME J 2010;4(9):1136–1143.

80. Chater KF. Recent advances in understanding Streptomyces. F1000Res 2016;5:2795.

81. Kim JN, Kim Y, Jeong Y, Roe JH, Kim BG et al. Comparative Genomics Reveals the Core and Accessory Genomes of Streptomyces Species. J Microbiol Biotechnol 2015;25(10):1599–1605.

82. Lioy VS, Lorenzi JN, Najah S, Poinsignon T, Leh H et al. Dynamics of the compartmentalized Streptomyces chromosome during metabolic differentiation. Nat Commun 2021;12(1):5221.

83. Szafran MJ, Małecki T, Strzałka A, Pawlikiewicz K, Duława J et al. Spatial rearrangement of the Streptomyces venezuelae linear chromosome during sporogenic development. Nat Commun 2021;12(1):5222.

84. Choudoir MJ, Buckley DH. Phylogenetic conservatism of thermal traits explains dispersal limitation and genomic differentiation of Streptomyces sister-taxa. ISME J 2018;12(9):2176–2186.

85. Andam CP, Choudoir MJ, Vinh Nguyen A, Sol Park H, Buckley DH. Contributions of ancestral inter-species recombination to the genetic diversity of extant Streptomyces lineages. ISME J 2016;10(7):1731–1741.

86. Tidjani AR, Lorenzi JN, Toussaint M, van Dijk E, Naquin D et al. Massive Gene Flux Drives Genome Diversity between Sympatric. MBio 2019;10(5).

87. McDonald BR, Currie CR. Lateral Gene Transfer Dynamics in the Ancient Bacterial Genus. Mbio 2017;8(3).

88. Kun Cheng XR, Ying Huang,. Widespread interspecies homologous recombination reveals reticulate evolution within the genus Streptomyces. Molecular Phylogenetics and Evolution; 2016; 102:246–254

89. Zhang Z, Shitut S, Claushuis B, Claessen D, Rozen DE. Mutational meltdown of putative microbial altruists in Streptomyces coelicolor colonies. Nat Commun 2022;13(1):2266.

90. Chung Y-H, Kim H, Ji C-H, Je H-W, Lee D et al. Comparative Genomics Reveals a Remarkable Biosynthetic Potential of the Streptomyces Phylogenetic Lineage Associated with Rugose-Ornamented Spores. Msystems 2021;6(4):e00489–00421.

